## Supplementary Figures for "Non-coding AUG circRNAs constitute an abundant and conserved subclass of circles"

**Supplementary Figures 1-20**

#### Supplementary Figure 1

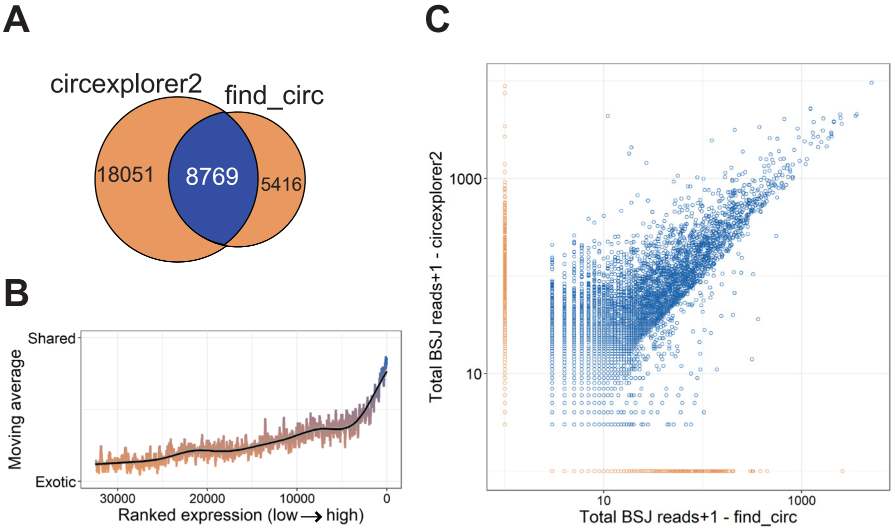


**Supplementary Figure 1: CircRNAs prediction in ENCODE data from mouse (Relates to Fig. 1A-C). A**) Venndiagram showing the number of exotic (orange) and shared (blue) circRNAs found by find_circ and circexplorer2 algorithms in the ENCODE datasets from mouse. **B**) Smoothed fraction of shared circRNAs found by find_circ and circexplorer2 as a function of ranked expression. **C**) Scatterplot depicting the number of backsplice junction (BSJ)-spanning reads obtained from find_circ and circexplorer2 across all the samples analysed. The points are color-coded as shared (blue) or exotic (orange).

### Supplementary Figure 2

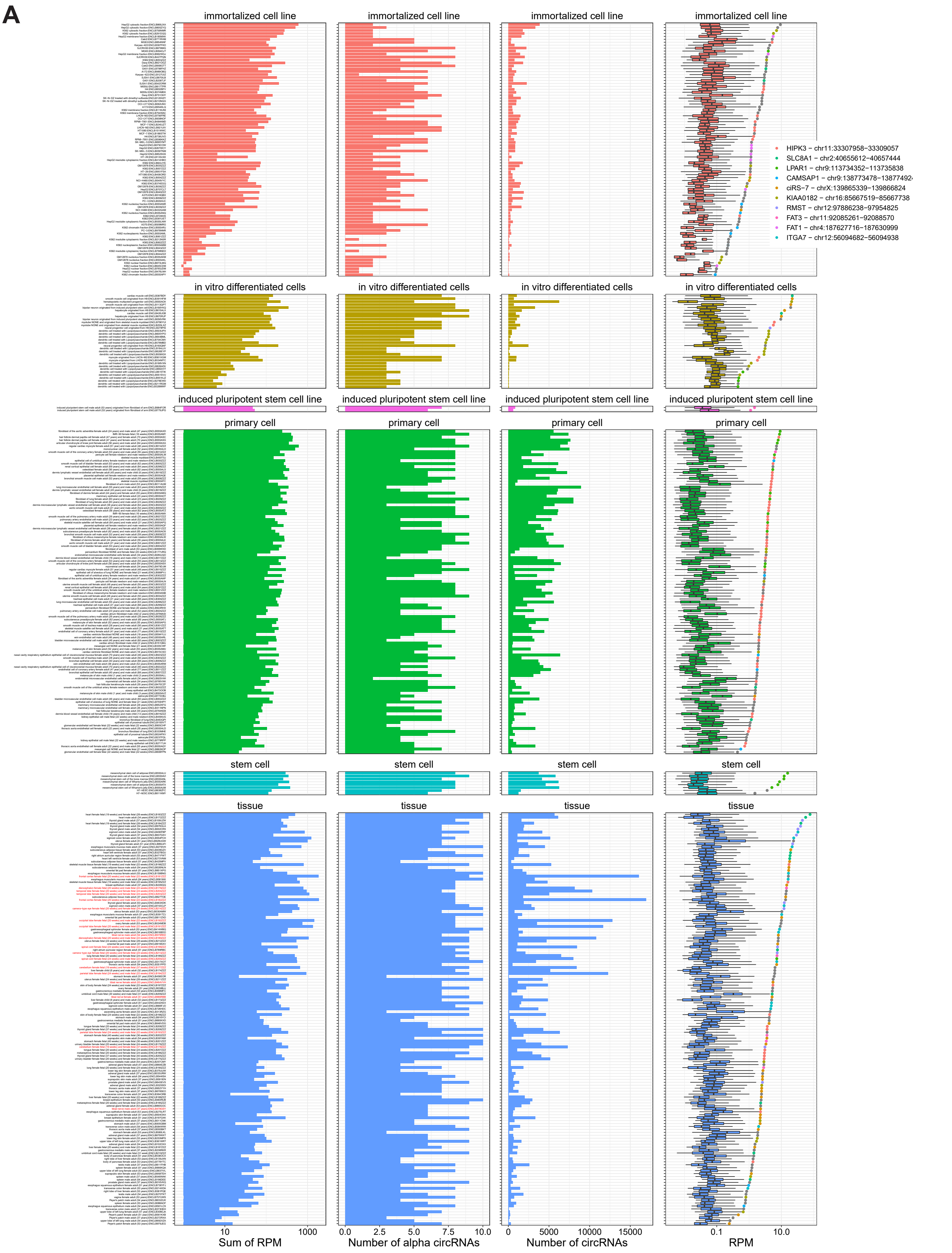


**Supplementary Figure 2: CircRNA and mRNA expression in each sample (3 pages). A-D**) For all human (A-B) and mouse (C-D) samples analyzed, the circRNA (A,C) and mRNA (B,D) expression is shown. The samples are subgrouped into biosamples as annotated by ENCODE and arranged by alpha expression. The panels show overall circRNA or mRNA content (1st panel), number of unique circRNAs or mRNAs identified (2$^{nd}$ panel), number of top10 alpha circR-

#### Supplementary Figure 2 (continued)

**B**

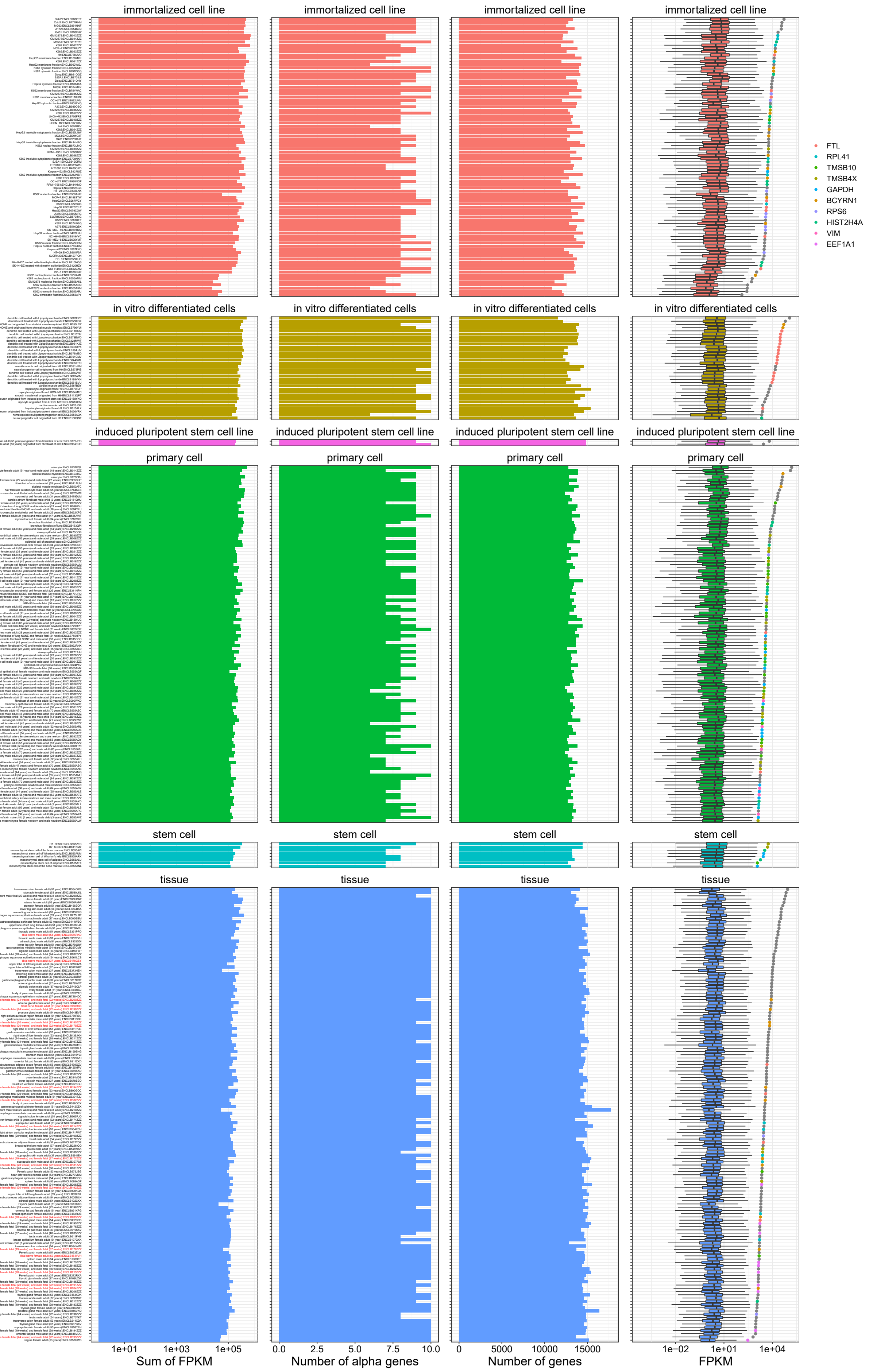


NAs/mRNAs identified (3rd panel) and finally the distribution of expression depicted as boxplot (last panel) including alpha circRNA/mRNA expression (color-coded as shown in legend; grey ~ alpha circRNA/mRNA not in top10). Brain-derived samples are highlighted in red to the left.

### Supplementary Figure 2 (continued)

C

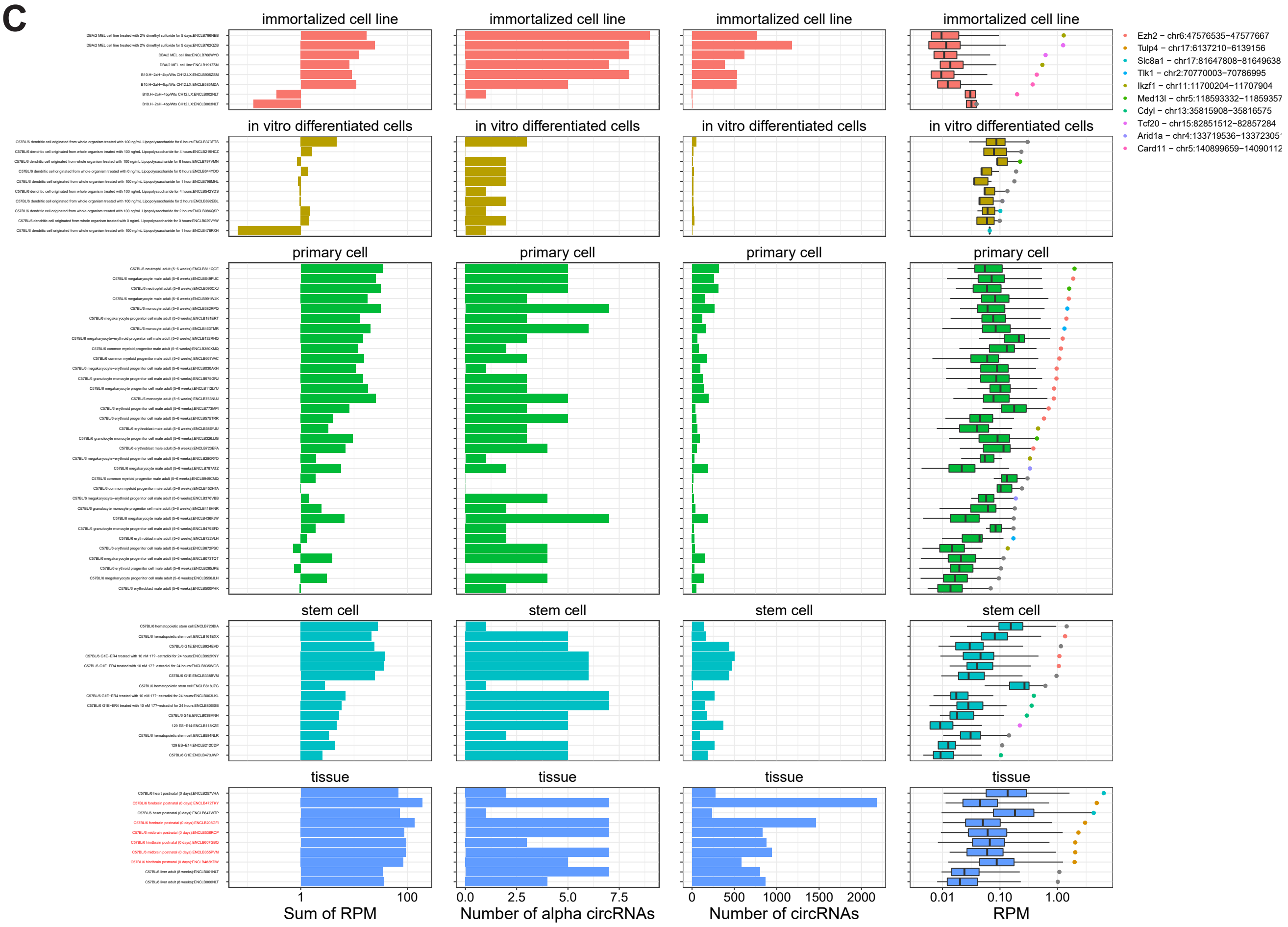


D

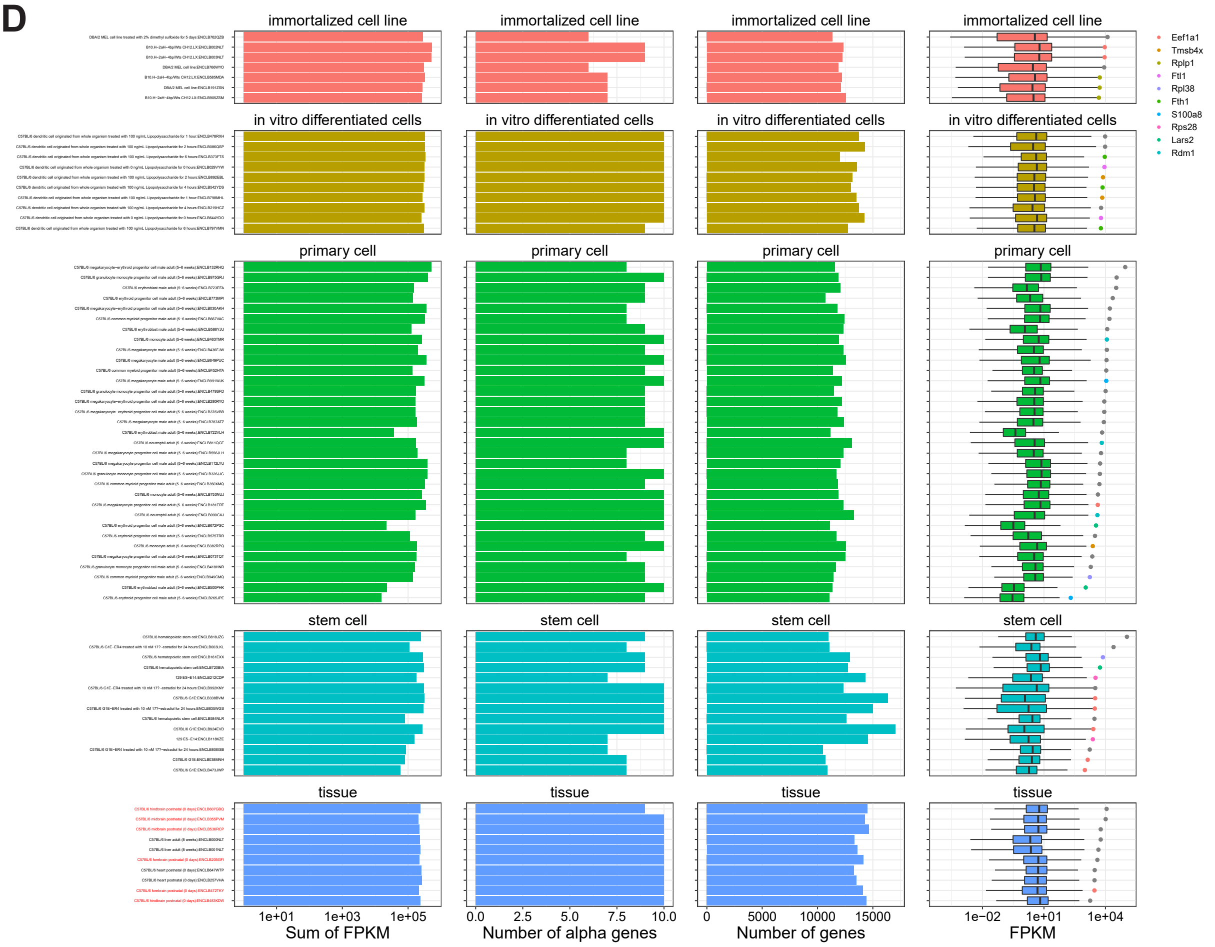

#### Supplementary Figure 3

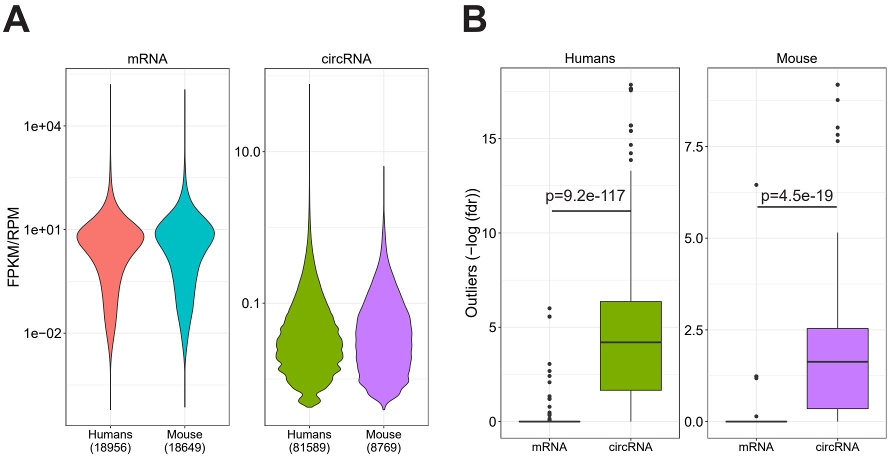


**Supplementary Figure 3: Expression and outlier analysis. A**) Violin plot of mRNA FPKM values across all samples for human and mouse as denoted, or RPM values for all circRNAs across samples for human and mouse. The number of unique mRNAs and circRNAs detected is denoted in parenthesis. **B**) Boxplot depicting the one-tailed Grubbs-test-based p-values for alpha mRNA and alpha circRNA outlier expression in humans and mice as denoted. P-values are adjusted for multiple testing using the Benjamini-Hochberg procedure (fdr). Differences between mRNAs and circRNAs fdr-distributions are evaluated with Wilcoxon rank-sum test.

#### Supplementary Figure 4

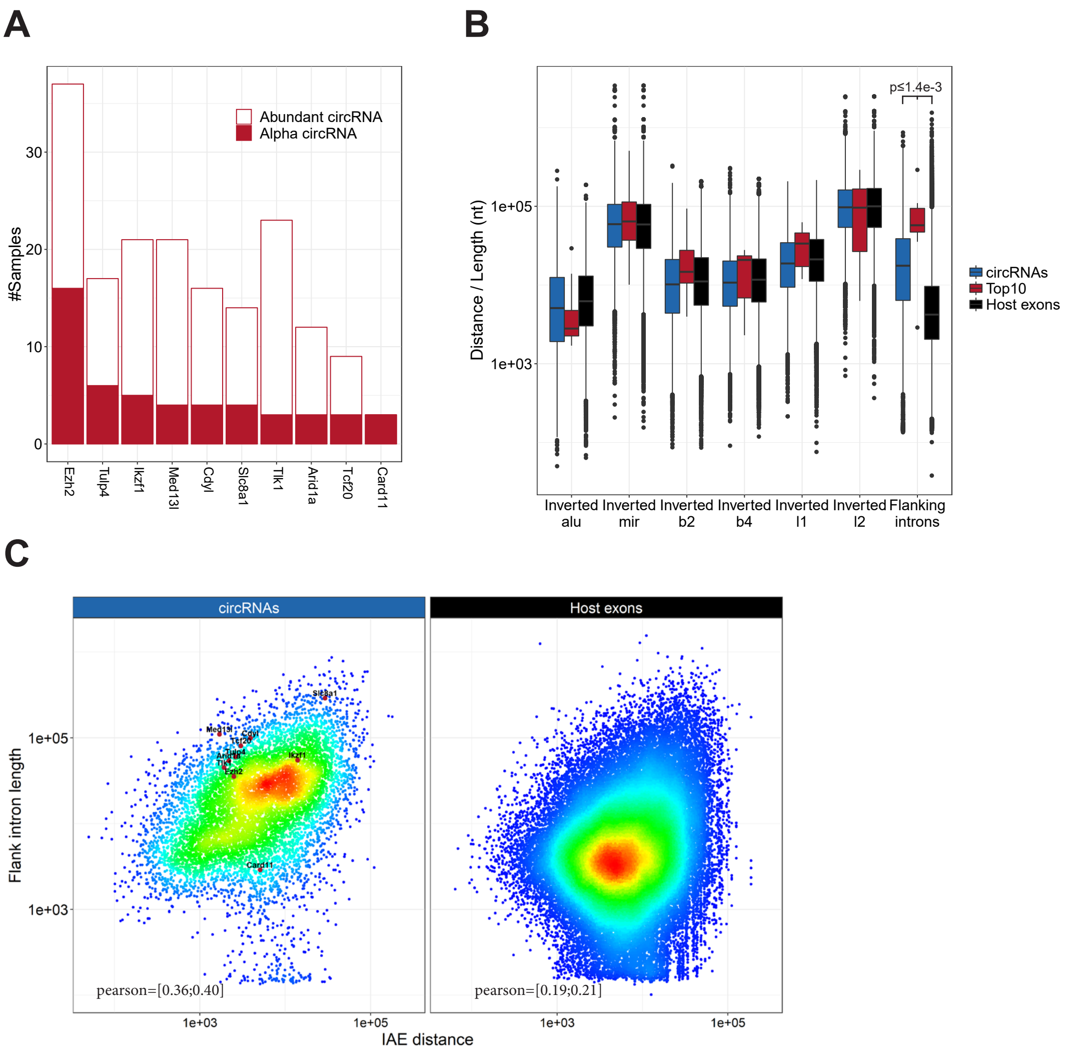


**Supplementary Figure 4: Alpha circRNAs (Relates to Fig. 1D-F). A**) The alpha frequency of the top 10 most commonly found alpha circRNAs in the murine samples as well as the frequency of being an abundant circRNA (i.e. within the top 10 expressed circRNAs in a sample) are plotted as a stacked barplot. **B**) Boxplot comparing the distance to inverted repeat element and flanking intron length for circRNAs in general, host gene exons and the top 10 murine alpha circRNAs (see schematics in Fig. 1D). **C**) Density-colored scatterplot showing relationship between IAE (using B1/alu elements) and flanking intron length for all circRNAs (left) and host gene exons (right). The top 10 murine alpha circRNAs are highlighted to the left.

#### Supplementary Figure 5

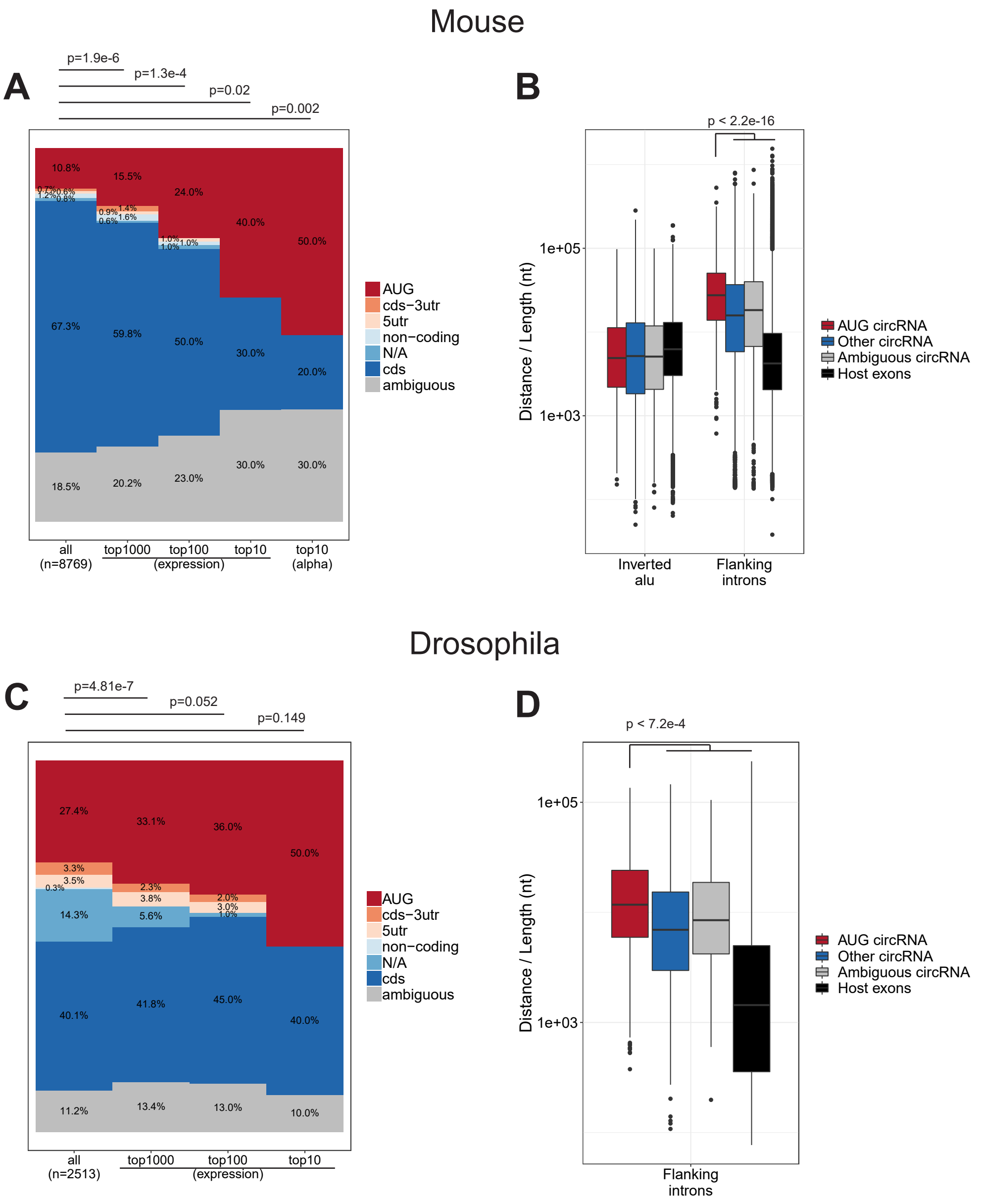


**Supplementary Figure 5: AUG circRNA in mouse and drosophila (Relates to Fig. 2B and E). A-B**) Frequency of circRNA- annotations for either all circRNA, the top 1000, top 100, top 10 circRNAs in mouse ENCODE data (A) or drosophila circRNAs found by Westholm et al, 2014 (B) based on overall expression (BSJ-spanning reads) color-coded as denoted. In (A) the annotation-distribution for the top 10 murine alpha circRNAs is included.  P-values are calculated using Fisher's exact test. **C-D**) Boxplot depicting the AIE distance (using B1/alu elements) and flanking intron length for all murine circRNAs (stratified by annotation) and host gene exons (C), or boxplot on flanking intron length for all drosophila circRNAs from Westholm et al, 2014, stratified by annotation. P-value is based on Wilcoxon rank-sum tests between 'AUG circRNA' and any of the other subgroups – only the highest p-value is shown.

#### Supplementary Figure 6

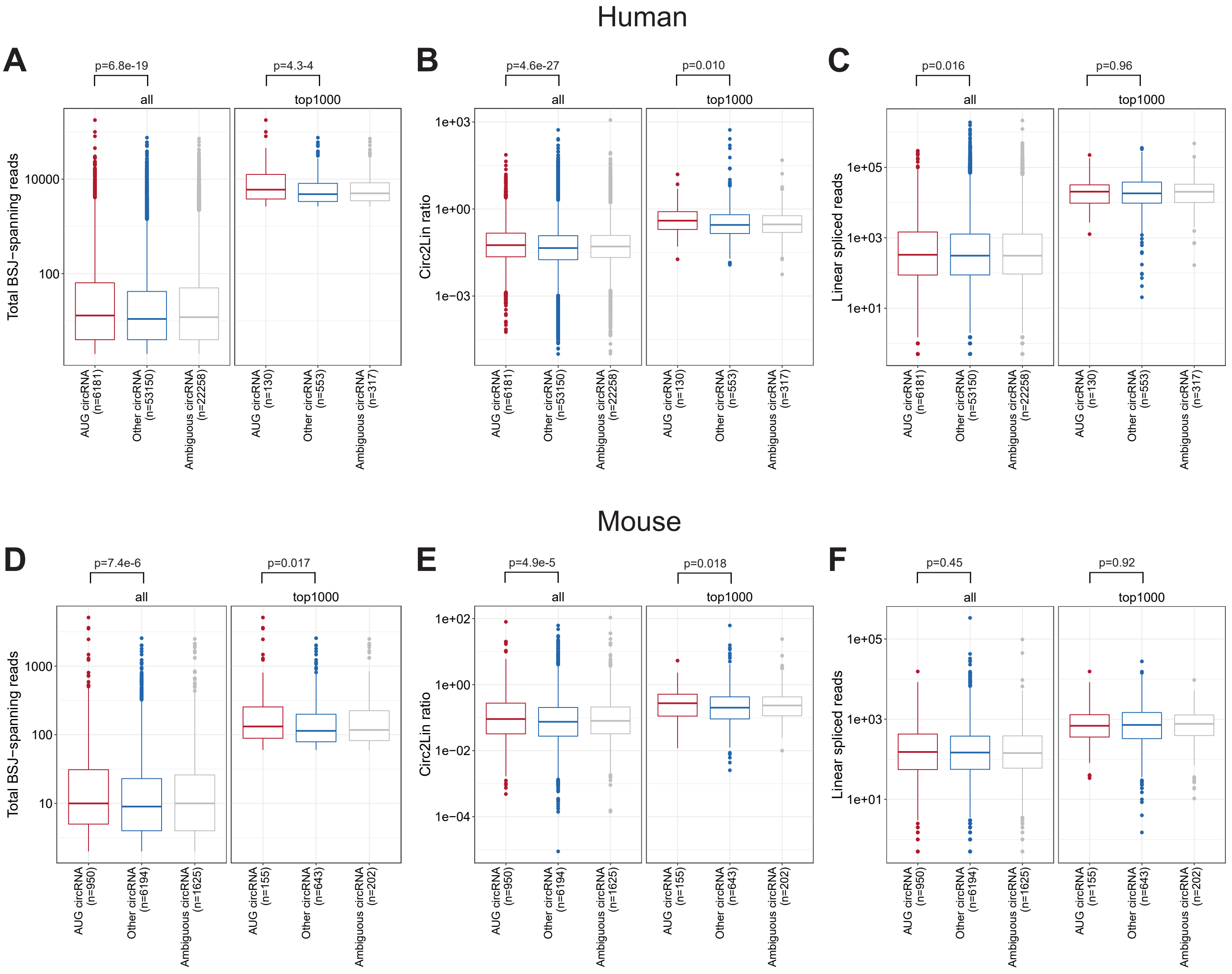


**Supplementary Figure 6: AUG circRNA expression. A-C**) Total BSJ-spanning reads (A), circular-to-linear ratios, (B), or linear spliced reads (C) for 'AUG circRNA', 'Ambiguous circRNA' and 'Other circRNAs' overall or within the top 1000 expressed human circRNAs. **D-F**) As in (A-C) but instead for murine circRNAs. P-values are based on Wilcoxon rank-sum test.

#### Supplementary Figure 7

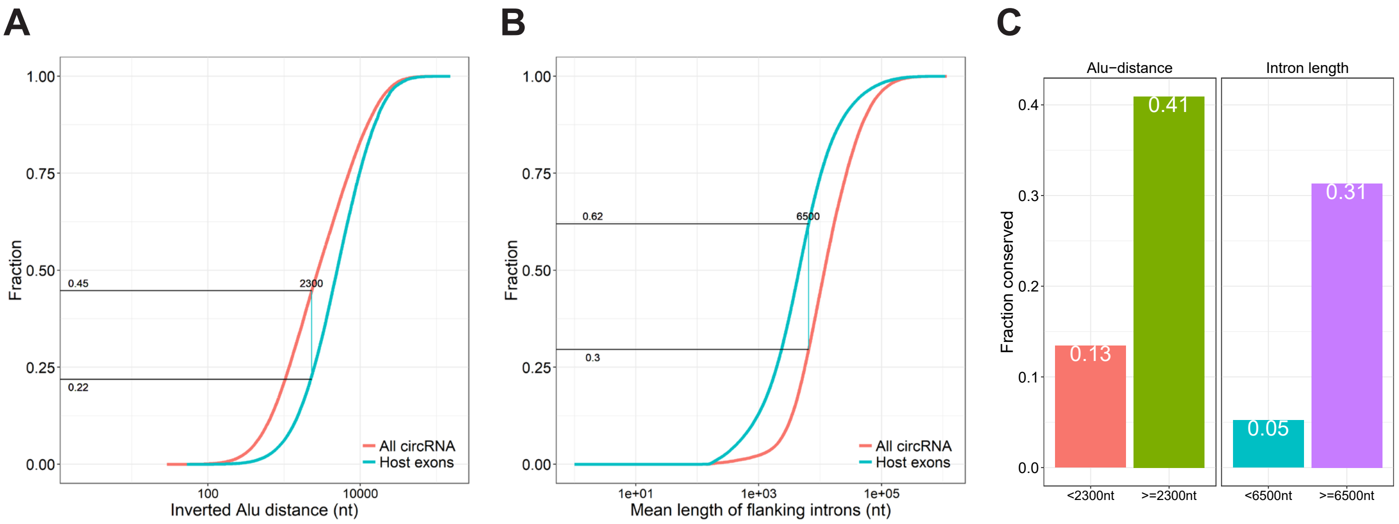


**Supplementary Figure 7: Cutoffs Inverted *Alu* element distance and flanking intron length. A-B**) Cumulative plot of inverted Alu element (IAE) distance (A) and flanking intron lengths (B) for all human circRNAs (n=81589) and corresponding host-gene exons (n=131002). The empirical cutoff, i.e. value corresponding to highest difference between circRNA and host gene exons, is denoted and the associated fraction for circRNA and host exons below the cutoff, respectively. **C**) Within the top 1000 expressed human circRNAs, the fraction of conserved circRNA above or below the AIE distance and flanking intron length cutoffs.

#### Supplementary Figure 8

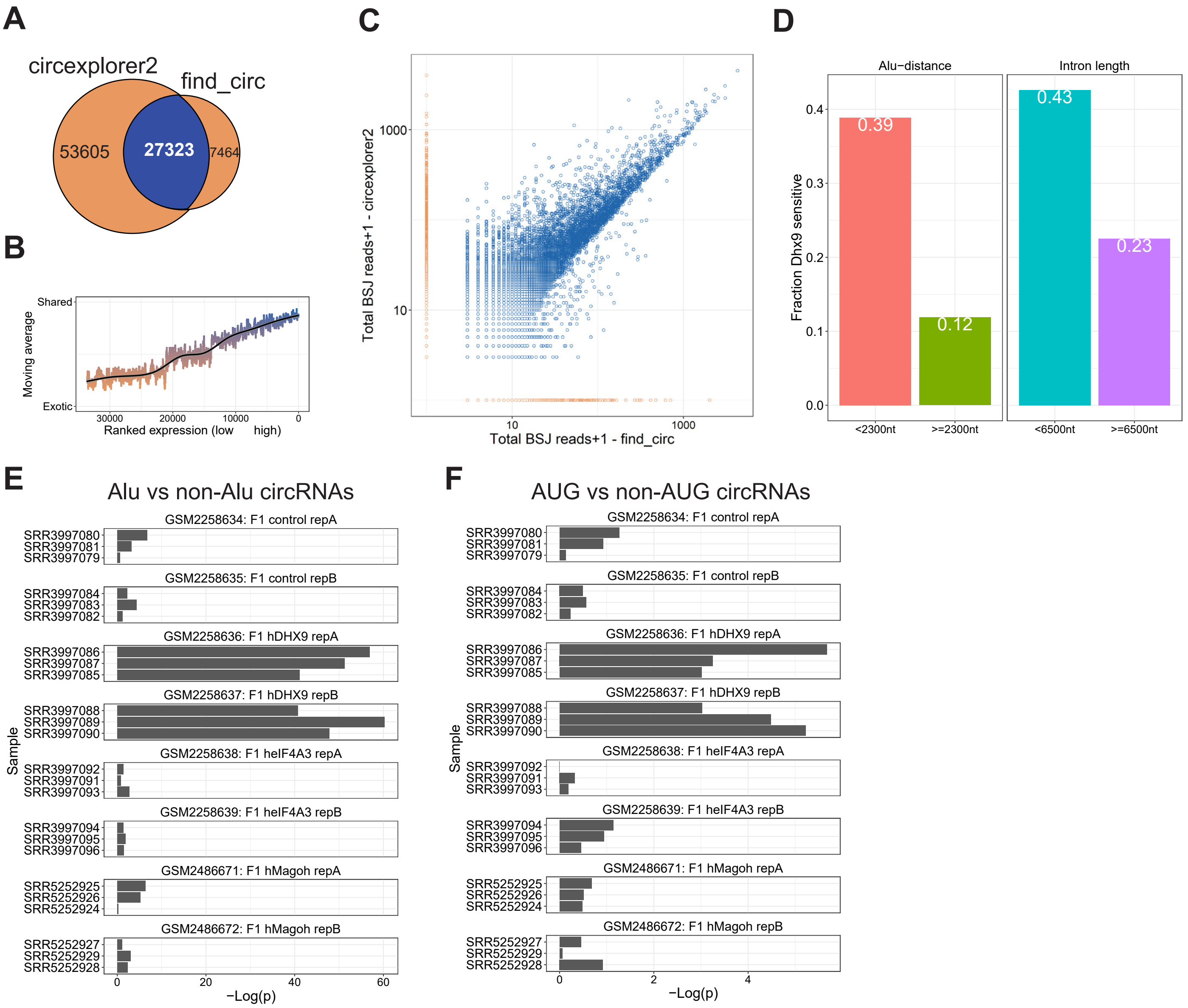


**Supplementary Figure 8: Dhx9-sensitive circRNA (Relates to Fig. 3). A**) Venndiagram showing the number of exotic (orange) and shared (blue) circRNAs found by find_circ and CIRCexplorer2 algorithms in GSE85161 RNAseq. **B**) Smoothed fraction of shared circRNAs found by find_circ and circexplorer2 as a function of ranked expression. **C**) Scatterplot depicting the number of backsplice junction (BSJ)-spanning reads obtained from find_circ and CIRCexplorer2. The points are color-coded as shared (blue) or exotic (orange). **D**) Barplot depicting fraction of Dhx9-sensitive circRNAs, i.e. fdr < 0.05, with proximal or distal IAE (left) or with short or long flanking introns (right) using cutoffs as determined in Supplementary Fig. 7. **E-F**) CircRNAs were stratified by long/short IAE distance (E) or by AUG/non-AUG annotation (F), and the number of HITS-CLIP reads in the immediate flanking regions were retrieved. The –log (p)-values determined by Wilcoxon rank-sum tests are depicted for all HITS-CLIP datasets.

#### Supplementary Figure 9

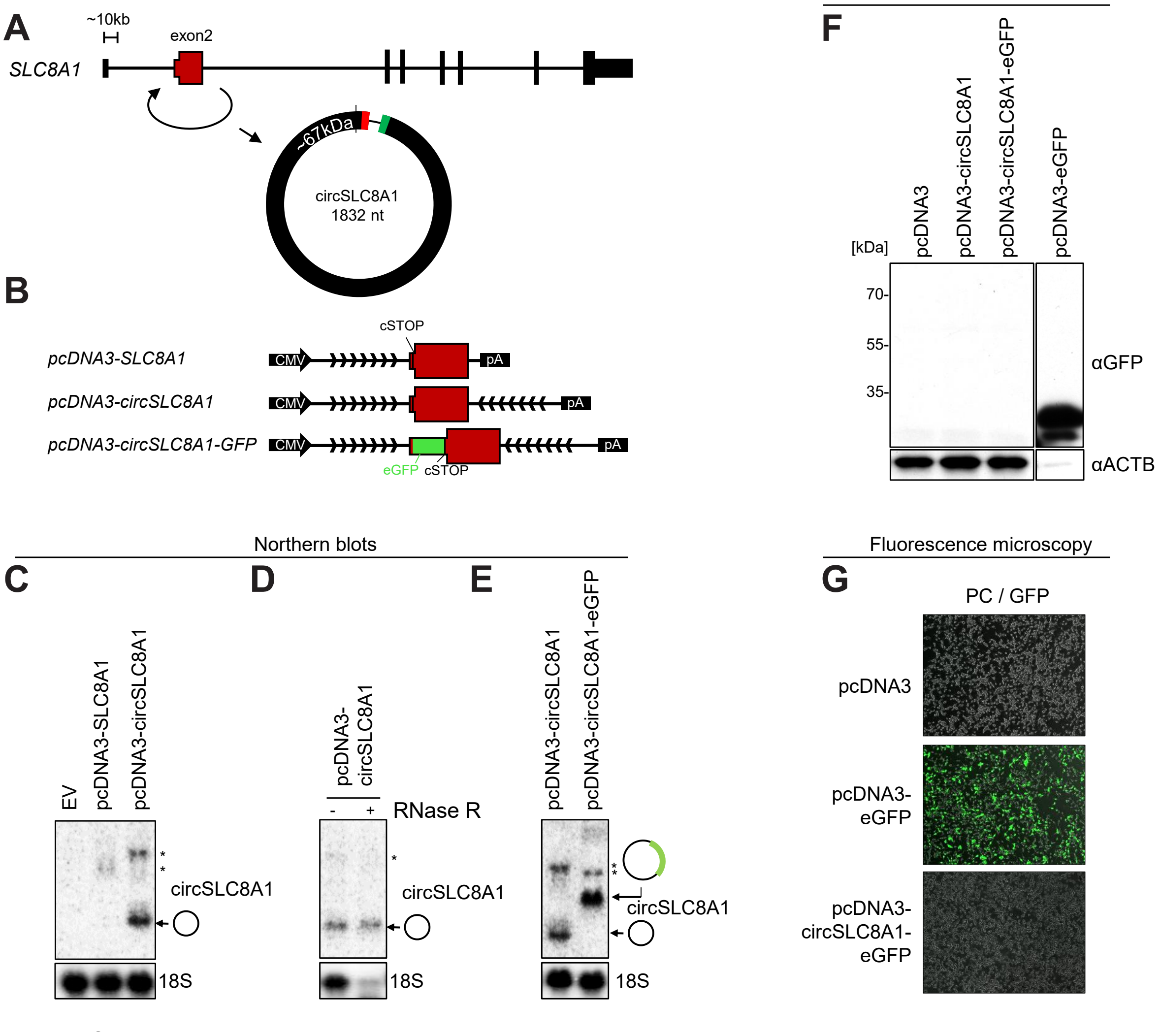


**Supplementary Figure 9: No evidence of circSLC8A1 translation. A**) Genomic representation of the *SLC8A1* hostgene locus. The exons are not drawn to scale. **B**) Schematic representation of expression vectors comprising the CMV promoter, the exon2 known to circularize, the putative circRNA-specific stop-codon (cSTOP), the insertion of eGFP ORF, the flanking regions (divergent arrows indicate artificially introduced inverted element) and the BGH pA signal. **C-E**) Northern blots showing effective expression of circRNA when flanked by inverted element (C), RNase R resistance (D) and circRNA production from the circSLC8A1-GFP fusion (E). Asterisks indicate non-circular by-products of the ectopic expression vectors. **F**) Western blot showing GFP expression from positive control (pcDNA-eGFP) and circSLC8A1-eGFP fusion. **G**) Merged phase contrast and GFP fluorescence images (PC/GFP) obtained from HEK293 cells transfected with vectors as denoted.

#### Supplementary Figure 10

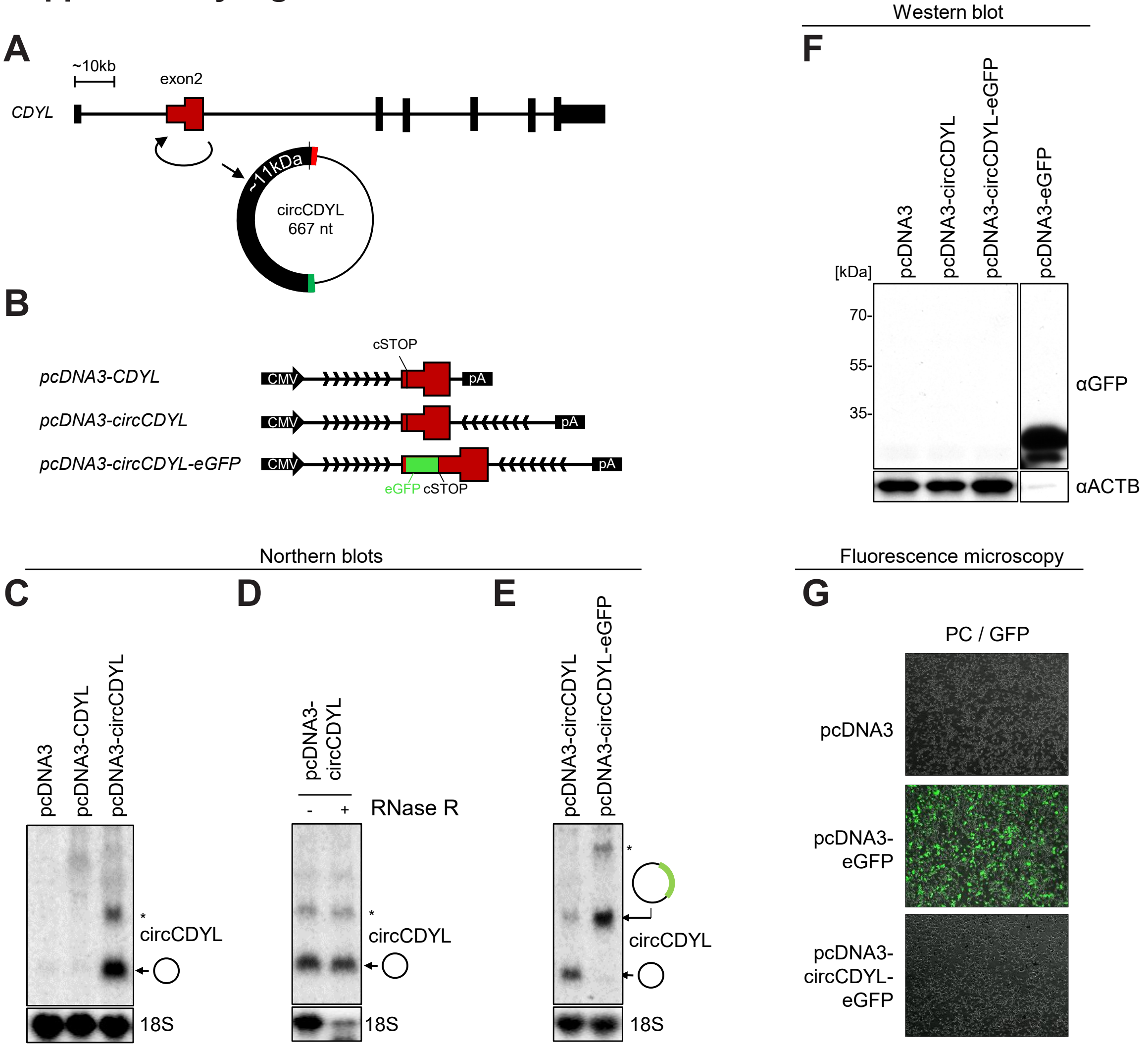


**Supplementary Figure 10: No evidence of circCDYL translation. A-G**) As in Supplementary Fig. 9, but using circCDYL expression vectors (see schematics in B). The western blot analysis (F) is from the same experiment as in **Supplementary Fig. 9F**, and the positive control lane (pcDNA3-eGFP) is the same.

#### Supplementary Figure 11

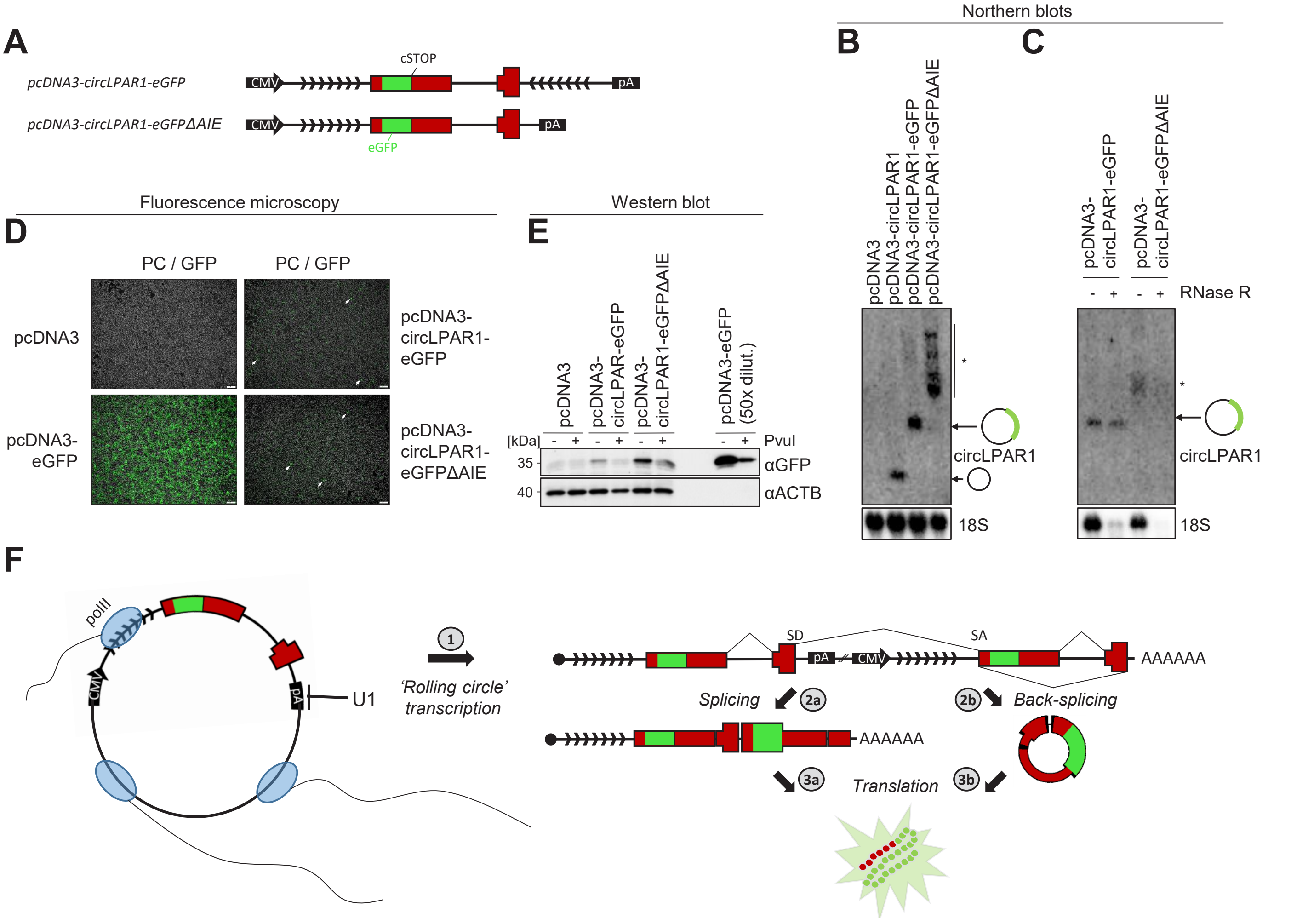


**Supplementary Figure 11: BSJ production from exon-repeat artefact. A**) Schematics showing circLPAR1-eGFP and circLPAR1-eGFPΔAIE expression vectors. **B-C**) Northern blot analysis on total RNA (B) or RNase R treated RNA (C) from circLPAR1-eGFP overexpression in HEK293T cells. 18S serves as loading and RNase R control. **D-E**) Merged phase contrast and GFP fluorescence images (PC/GFP) (D) and western blot analysis of GFP expression (E) from HEK293T cells transfected with vectors as denoted. **F**) Schematic model showing exon-repeat production as a consequence of 'rolling circle' plasmid transcription presumable facilitated by U1-mediated suppression of SD-downstream poly(A) signals (pA) (1). Then, the downstream splice donor (SD) couples to the upstream splice accepter (SA) in a conventi-onal linear splicing event producing an exon-junction sequence (2a) indistinguishable from backsplicing (2b), and as shown exon-repetition and circRNAs give rise to the exact same protein product if translated (3a and b).

#### Supplementary Figure 12

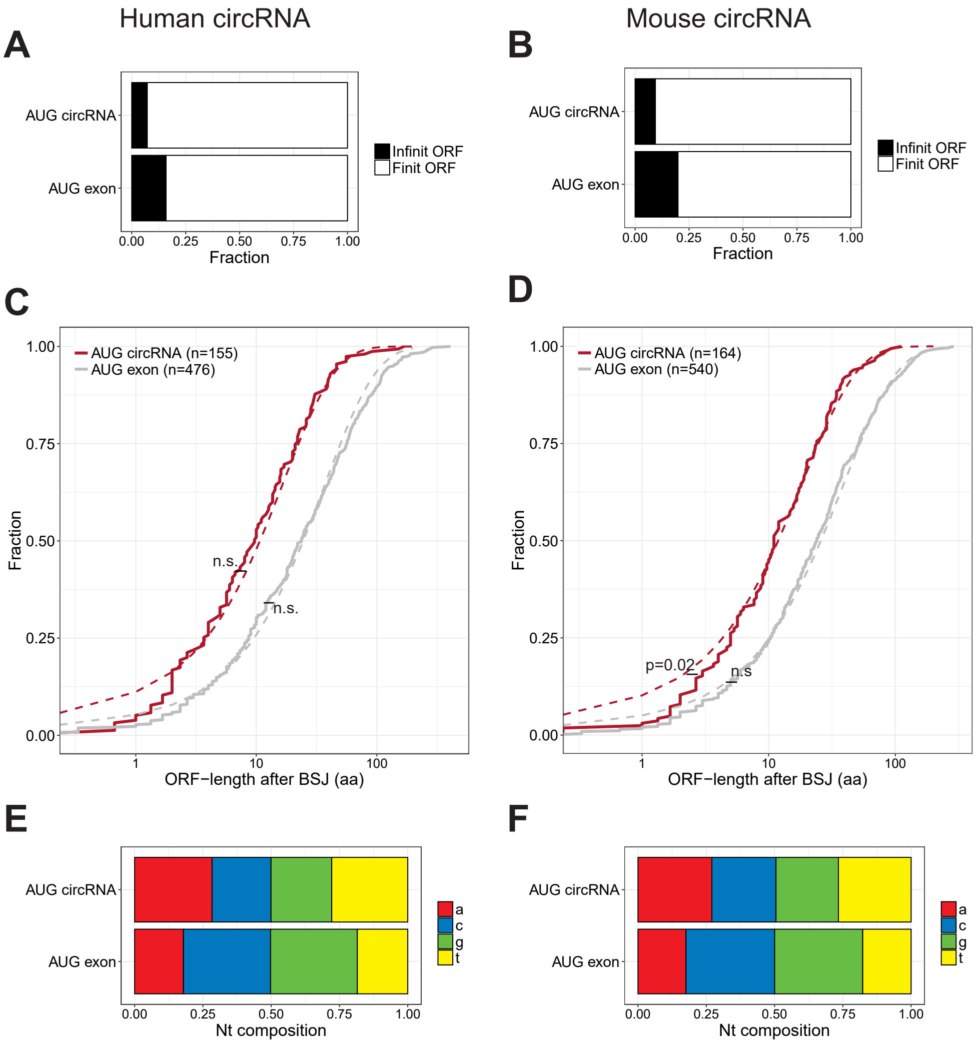


**Supplementary Figure 12: ORF analysis. A-B**) Fraction of predicted infinite ORFs in AUG circRNAs and AUG exons in humans (A) and mouse (B). **C-D**) Cumulative fraction plot of the distance from the BSJ to the predicted stop-codon in humans (C) and mouse (D) shown for both AUG circRNA and AUG exons. The dashed line corresponds to the geometric distribution of stop-codons considering the overall nucleotide composition (see below) in the 5'UTR of AUG circRNAs and AUG exons, respectively. P-values reflect chi-squared test on bins of different ORF lengths with at least five observations and the associated expected bin-size based on the geometric distribution (n.s.; not significant). **E-F**) Stacked barplot showing 5'UTR nucleotide composition of AUG circRNAs and AUG exons found in humans (E) and mouse (F).

#### Supplementary Figure 13

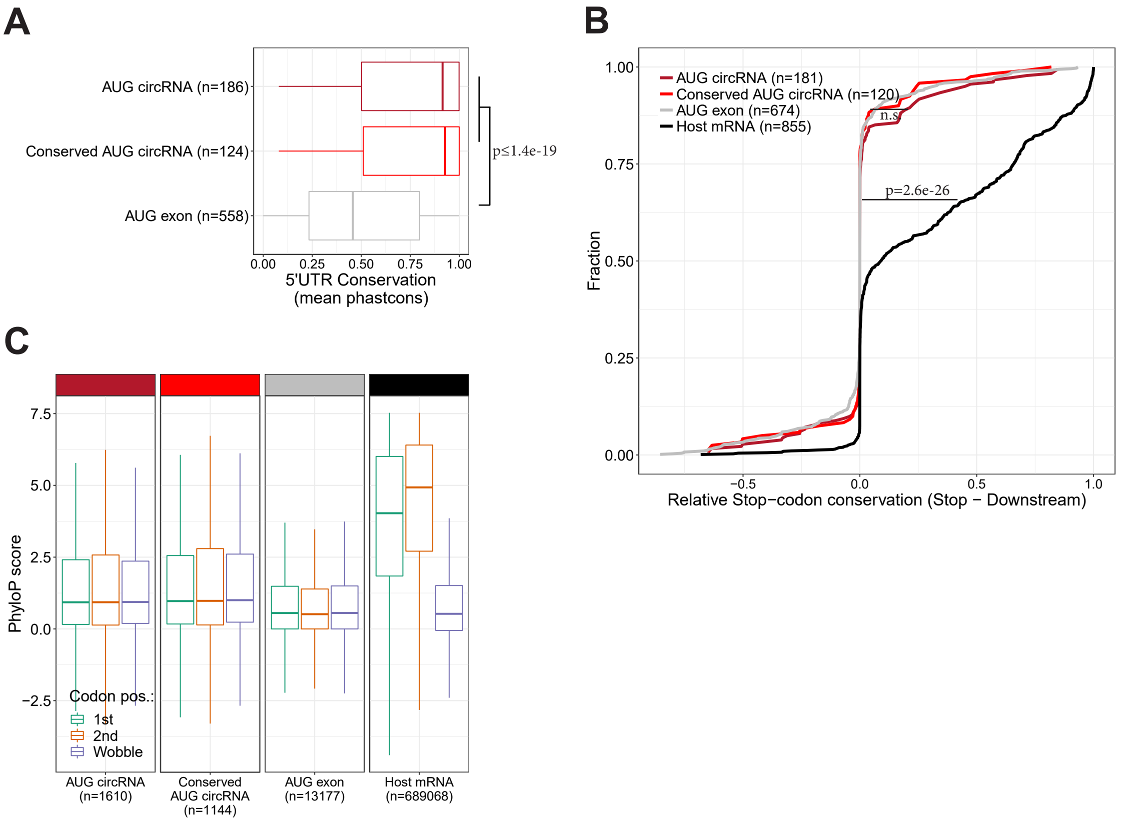


**Supplementary Figure 13: ORF conservation in mouse (relates to Fig. 5). A**) PhastCons analysis of 5'UTR within AUG-containing exons in all AUG circRNAs, conserved AUG circRNAs within top 1000 expressed murine circRNAs and as well as non-circular AUG exons from host-genes expression 'Other circRNAs' within top 1000. **B**) Analysis of stop codon conservation compared to immediately downstream triplet on putative circRNA-derived ORF (annotation as in (A)) as well as for bona-fide stop-codons within host-gene ORFs from mouse. P-value is determined by Wilcoxon rank-sum test. **C**) PhyloP analysis of single-position conservation for 1st, 2nd and wobble-position within putative ORF after BSJ and for bona-fide ORFs within host-genes from mouse.

### Supplementary Figure 14

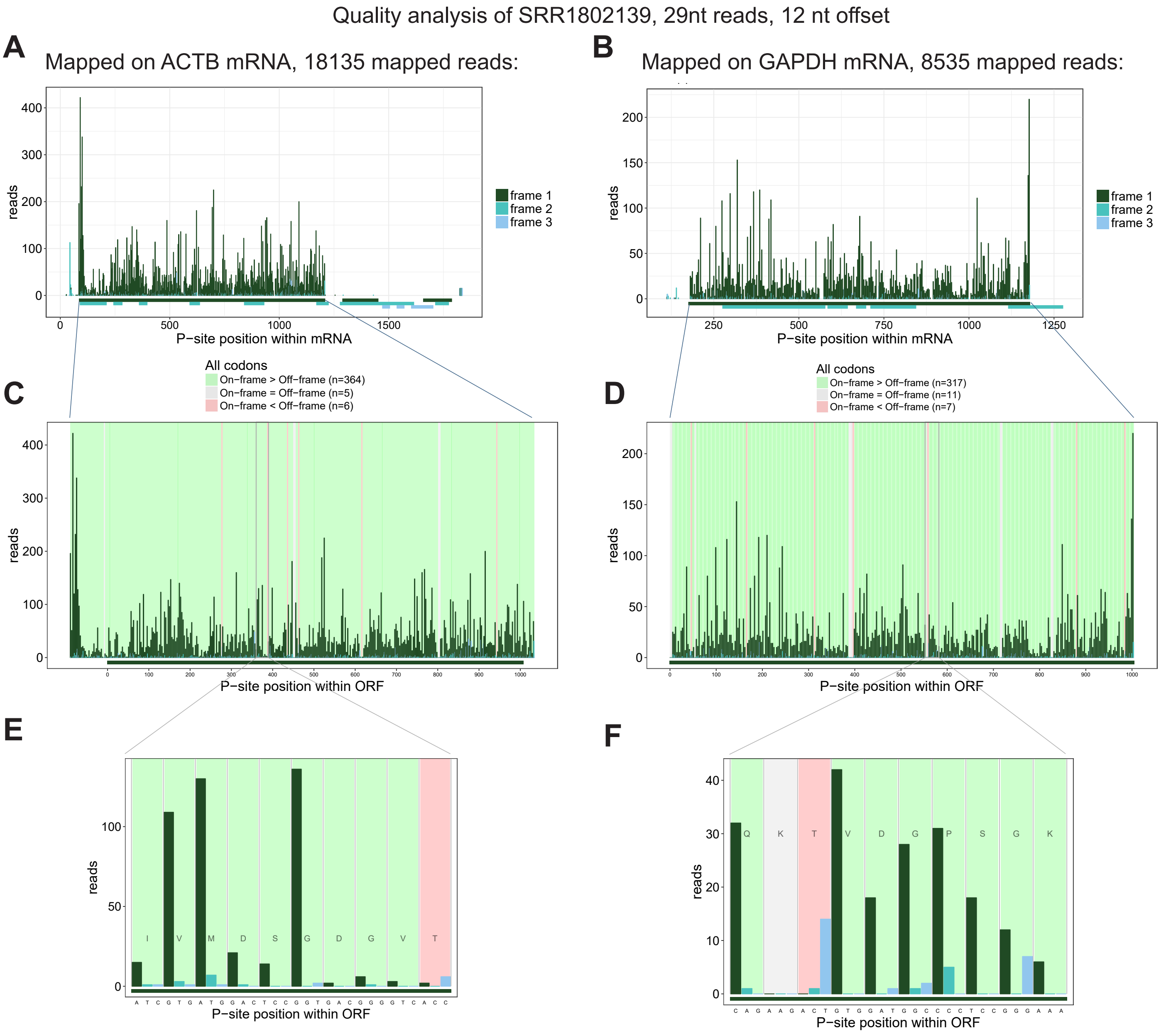


## G

p(ACTB)=binom.test (c(364,6), alternative=”greater”)= 1.44e-99
p(GAPDH)=binom.test (c(317,7), alternative=”greater”)= 2.08e-84
p(SRR1802139, 29nt readlength, 12nt offset)=(p(ACTB)+p(GAPDH))/2=1.04e-84

**Supplementary Figure 14: Example of Riboseq quality assessment. A-B**) 29nt Riboseq reads from SRR1802139 are mapped to *ACTB* mRNA (A) and *GAPDH* mRNA (B). P-site is here set at 12nt downstream the mapped 5’end. The number of reads as a function of P-site position is depicted for both mRNAs and color-coded by frame. All the predicted ORF with similar color-coding are shown below. **C-D**) Focusing specifically on the annotated ORF for *ACTB* (C) and *GAPDH* (D), the distribution of reads is shown (as in A-B). Moreover, for each codon, the background is color-coded according to the ratio between on-frame and off-frame reads, as denoted in the legend. **E-F**) Zooming in on a 10-codon stretch for both *ACTB* (E) and *GAPDH* (F) to show the phasing of reads and the corresponding background coloring. **G**) Calculation of p-values by binomial test. Here, for *ACTB*, 364 codons exhibit predominant on-frame reads, whereas seven codons have most off-frame reads (5 codons exhibit equal on and off-frame reads, but these are not considered in the statistics) resulting in P(*ACTB*)= 1.44e-99 calculated as shown. Similar procedure for *GAPDH* and the overall p-value for 29nt reads using 12nt offset from SRR1802139 is obtained from the mean of p(*ACTB*) and p(*GAPDH*).

#### Supplementary Figure 15

**A**

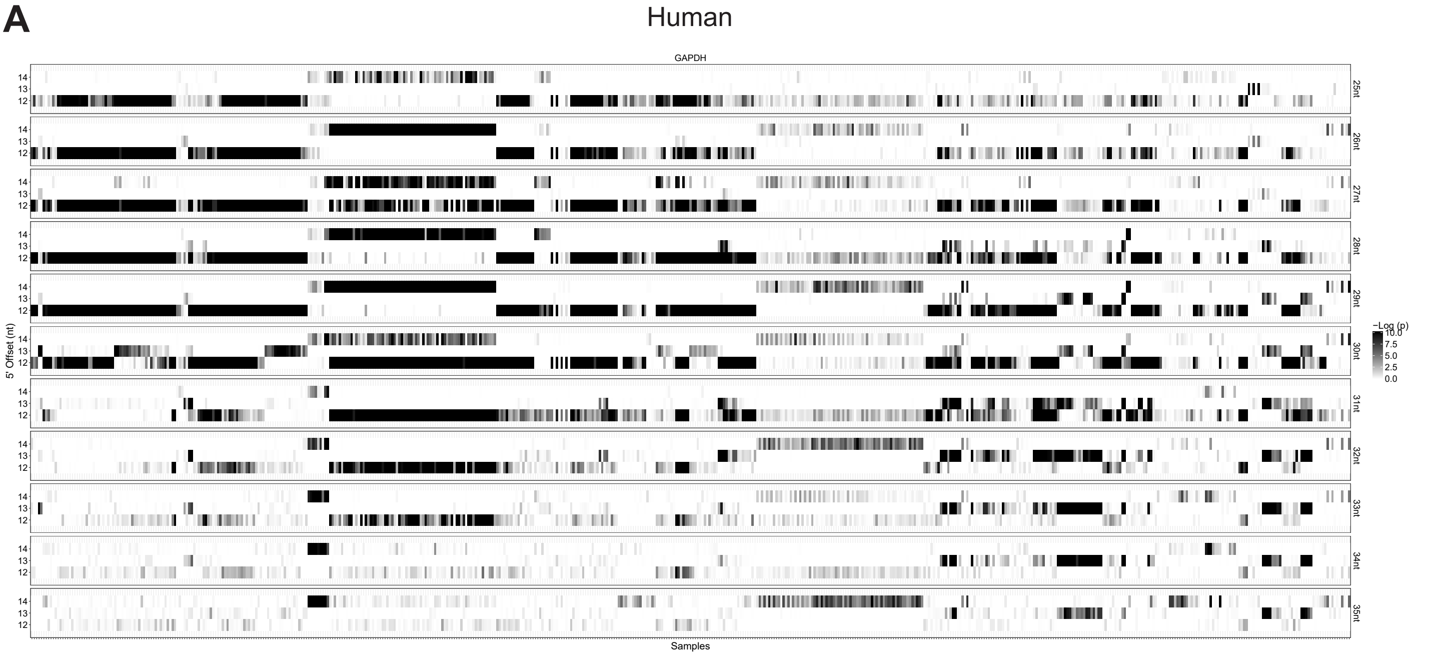


**B**

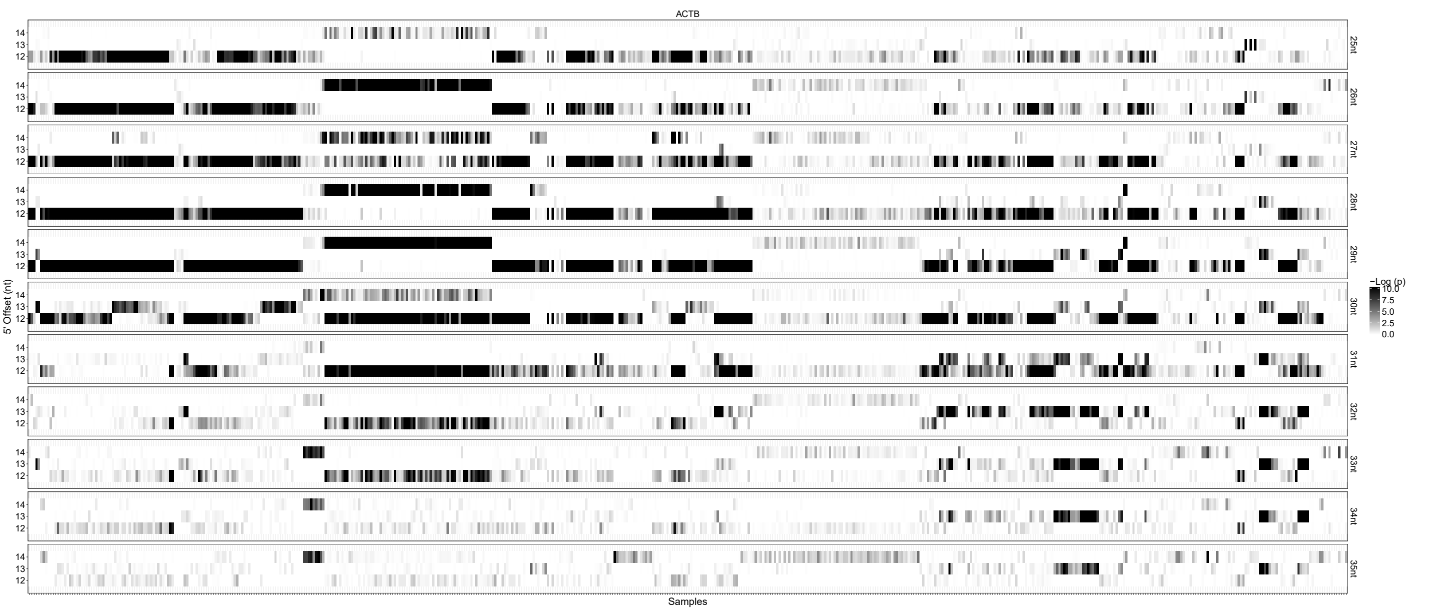


**Supplementary Figure 15: Riboseq quality assessment (2 pages)**. **A-D**) Heatmap of obtained –log(p) values for all riboseq samples (x-axis) analysing 25 nt to 35 nt reads individually (y-separated panels) using 12, 13, or 14 nt P-site offset (y-axis within panels) based on human samples (A-B) or mouse samples (C-D). The analyses were conducted on *ACTB* (A,C) and *GAPDH* mRNA (B,D), see Supplementary Fig. 11 for detailed description of quality assessment.

#### Supplementary Figure 15 (continued)

**C**

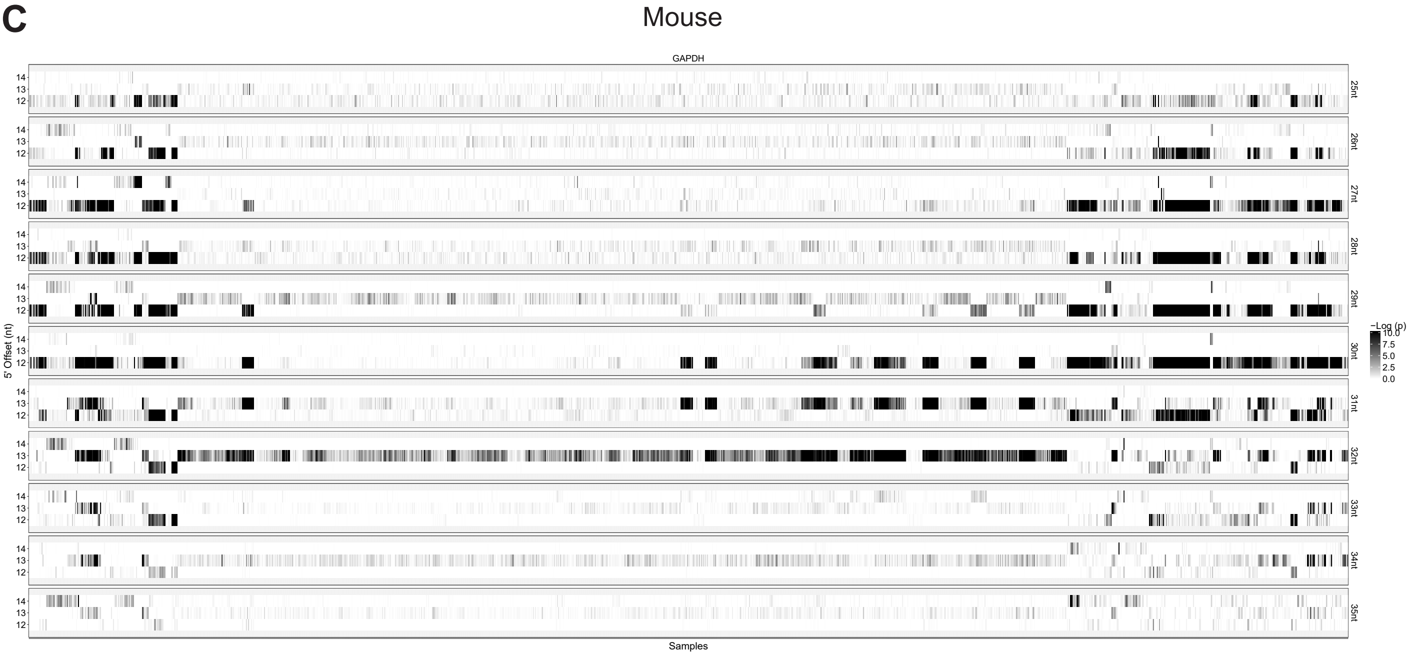


**D**

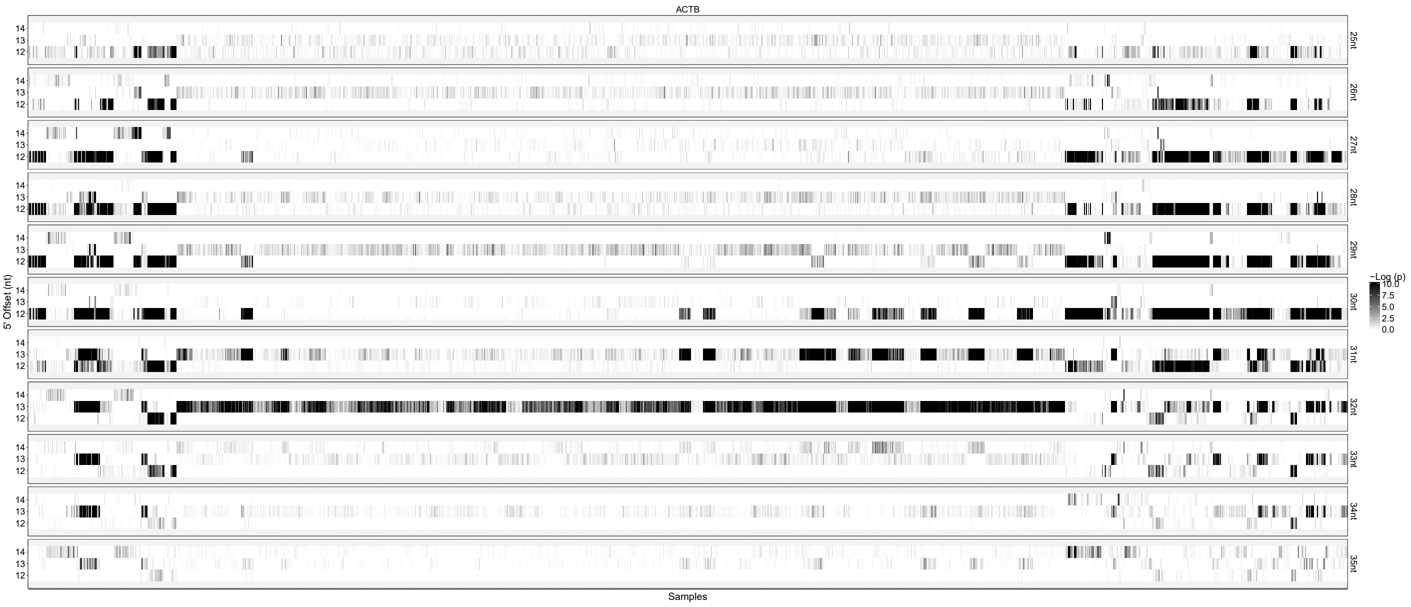

#### Supplementary Figure 16

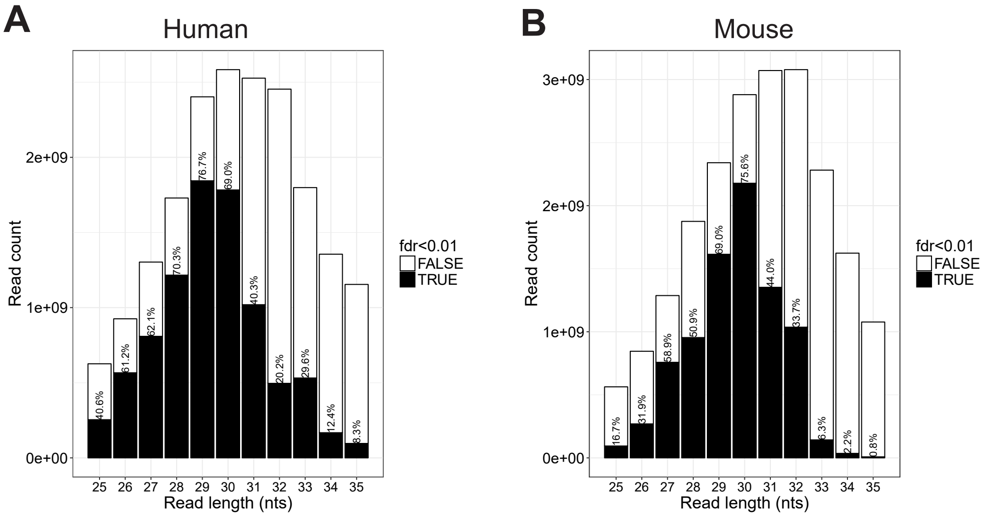


**Supplementary Figure 16: Riboseq quality and quantity by read-length overview. A-B**) For each read-length the number of high quality (fdr < 0.01) and low quality reads (fdr >= 0.01) are depicted for humans (A) and mouse (B) samples.

#### Supplementary Figure 17

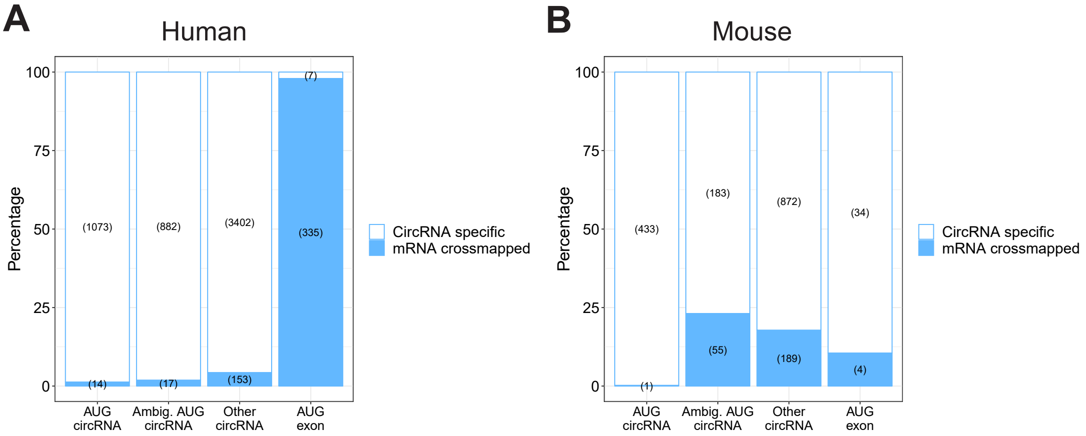


**Supplementary Figure 17: BSJ-derived reads mapping to mRNA transcriptome. A-B**) The fraction of RiboSeq reads derived from the BSJ grouped by circRNA type that also cross-map to the mRNA transcriptome with 1 mismatch tolerance in human (A) and mouse (B).

#### Supplementary Figure 18

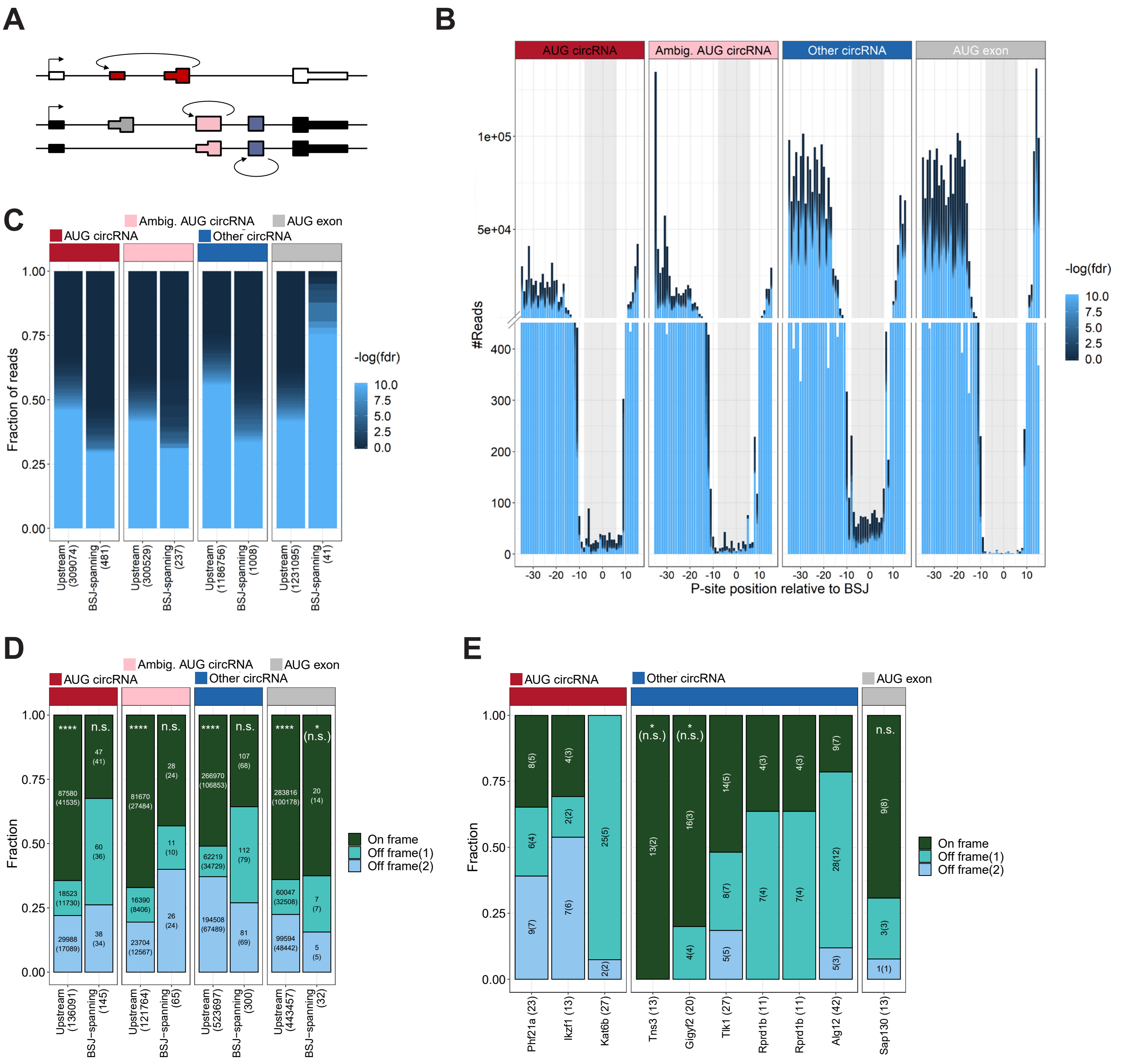


**Supplementary Figure 18: Ribosome profiling analysis across BSJ in mouse (Relates to Fig. 6). A**) Schematic overview showing circRNA annotation of 'AUG circRNA', 'Ambiguous circRNA', and 'Other circRNA'. **B**) Based on Ribosome profiling datasets, the number of ribosome P-sites across the backsplice junction (BSJ) was counted for each subclass of murine circRNAs (AUG circRNA, Ambiguous AUG circRNAs, and Other circRNA, see text for more detail), or the AUG-containing exon from non-AUG ('AUG exon') circRNA host-genes. The plot is color-scaled according to the associated read-class p-value (see Supplementary Fig. 15). The grey box denotes the defined P-site position of BSJ-spanning reads, from pos -8 to +6 relative to the BSJ. **C**) Based on all BSJ-spanning reads (-8 to +6) and upstream reads (-31 to -17), the read-class p-value distribution is shown. **D**) Based solely on reads classes with p<0.01, the fraction of P-sites in-frame and out-of-frame across the BSJ (-8 to +6) are shown for each subclass of circRNA. **E)** As in (D), but for each individual murine circRNA with 10+ reads across the BSJ.

#### Supplementary Figure 19

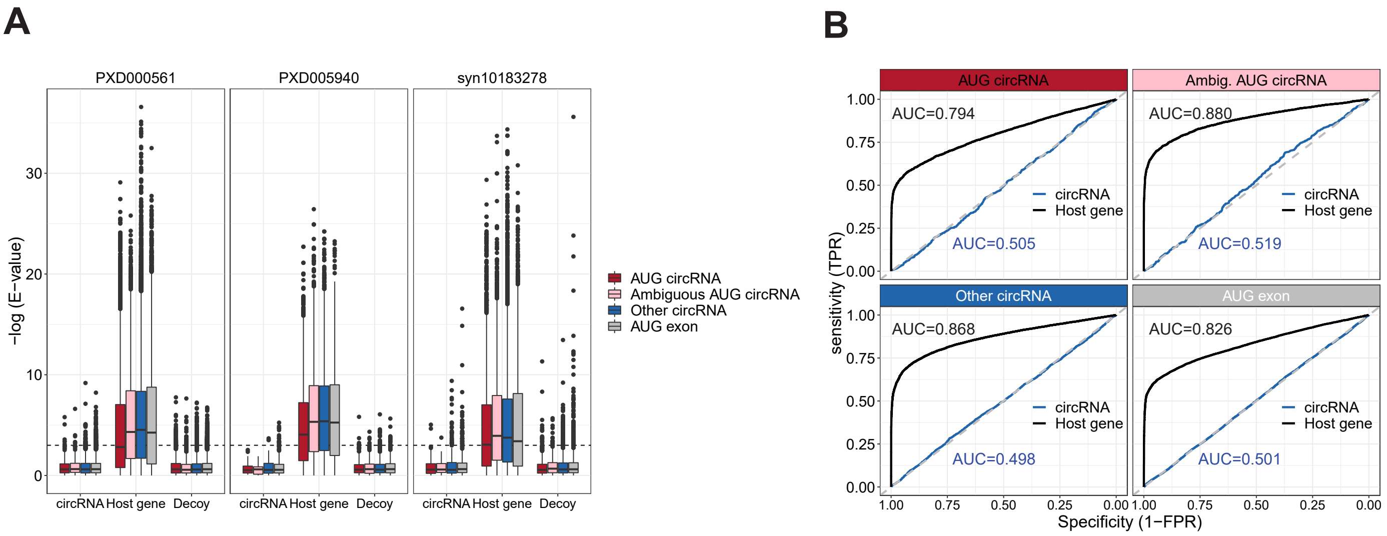


**Supplementary Figure 19: Mass-spec analysis of circRNA-derived peptides (Relates to Fig. 7). A)** Boxplot as in Fig. 7B but with each mass-spec accession analyzed separately. **B)** ROC curve on peptides identified as mRNA-derived or circRNA derived stratified by genic origin as denoted on strips. TPR (true positive rate) and FPR (false positive rate) are calculated as the cumulative sum of peptides divided by the total number of peptides, or the cumulative sum of decoy divided by the total number of decoys, respectively, on an e-value sorted list. Then, TPR is plotted as a function of 1-FPR as shown.

### Supplementary Figure 20

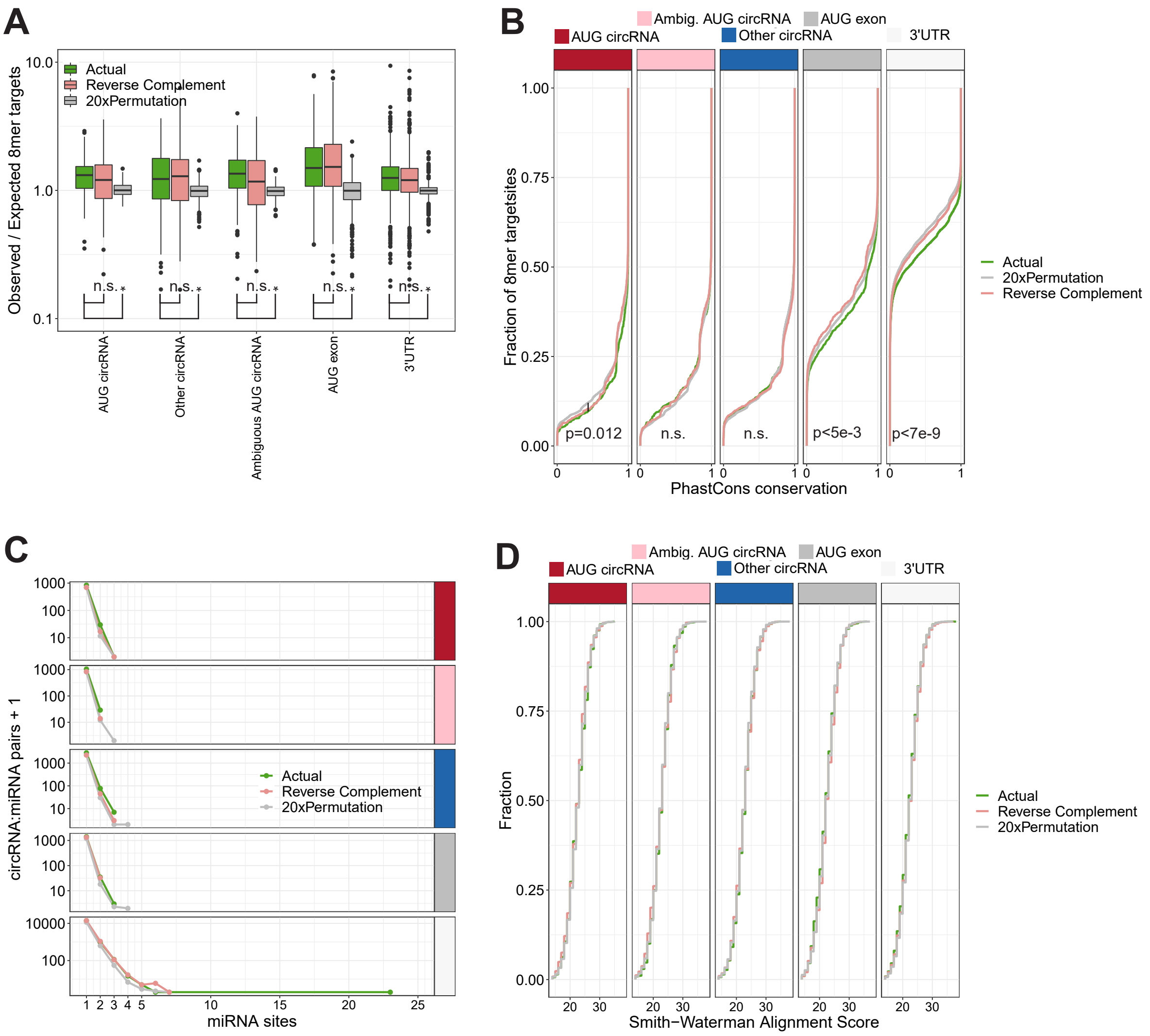


**Supplementary Figure 20: miRNA targets on circRNAs. A)** Total number of predicted 8mer target sites per circRNA, either the actual sequence, the reverse complement sequence and the mean output from 20x permutations (as per color-coding), normalized to the expected number of target sites considering the nucleotide composition of target sequence and target site. Only conserved and confident miRNAs from miRBase v21 was used (n=338). The target sequences are stratified by genic origin, and AUG exons and 3'UTR have been included in the analysis (x-axis). **B)** PhastCons conservation of the putative 8mer target sites observed in (A) depicted as a cumulative fraction plot. For the 20xPermutation, the phastCons scores reflect the conservation of the same position in the actual sequence. **C)** Frequency circRNA:miRNA pairs in the list of 8mer targetsites stratified genic origin. **D)** Smith-waterman alignment scores of all putative 8mer targetsites depicted as a cumulative fraction plot and stratified by genic origin. P-values are calculated by Wilcoxon rank-sum test (*, $p < 0.05$; n.s., not significant).
